# Overcoming Ligand Discovery Challenges: Developing Peptide-Based Tracers for SPSB2

**DOI:** 10.1101/2025.09.03.673904

**Authors:** Christopher Lenz, Lewis Elson, Johannes Dopfer, Frederic Farges, Andreas Krämer, Frank Löhr, Susanne Müller, Stéphanie M. Guéret, Herbert Waldmann, Volker Dötsch, Krishna Saxena, Stefan Knapp

## Abstract

Developing new E3 ligase ligands for the design of heterobivalent molecules, such as PROteolysis TArgeting Chimeras (PROTACs), requires careful evaluation of target engagement (TE). Characterizing protein-protein interactions (PPIs) is therefore essential in drug discovery, as it enables the assessment of ligand binding to sites that are often difficult to target. Degrons, peptide motifs recognized by E3 ligases, may serve as valuable starting points for designing E3 ligands. However, many degrons are highly polar and lack intrinsic membrane permeability, requiring alternative strategies for efficient cellular delivery. In this study, we used the SPRY domain-containing SOCS box protein 2 (SPSB2) E3 ligase as a model system to develop TE strategies for *in vitro* and *in cellulo* using polar degron-based peptides. By conjugating various polycationic cell-penetrating peptides (CPPs) to the degron sequence, we present a study demonstrating efficient cellular delivery. We obtained a high-resolution crystal structure and used various biophysical techniques to assess the influence of each modification, while confocal microscopy and BRET-based assays confirmed successful cellular delivery as well as potent target engagement.

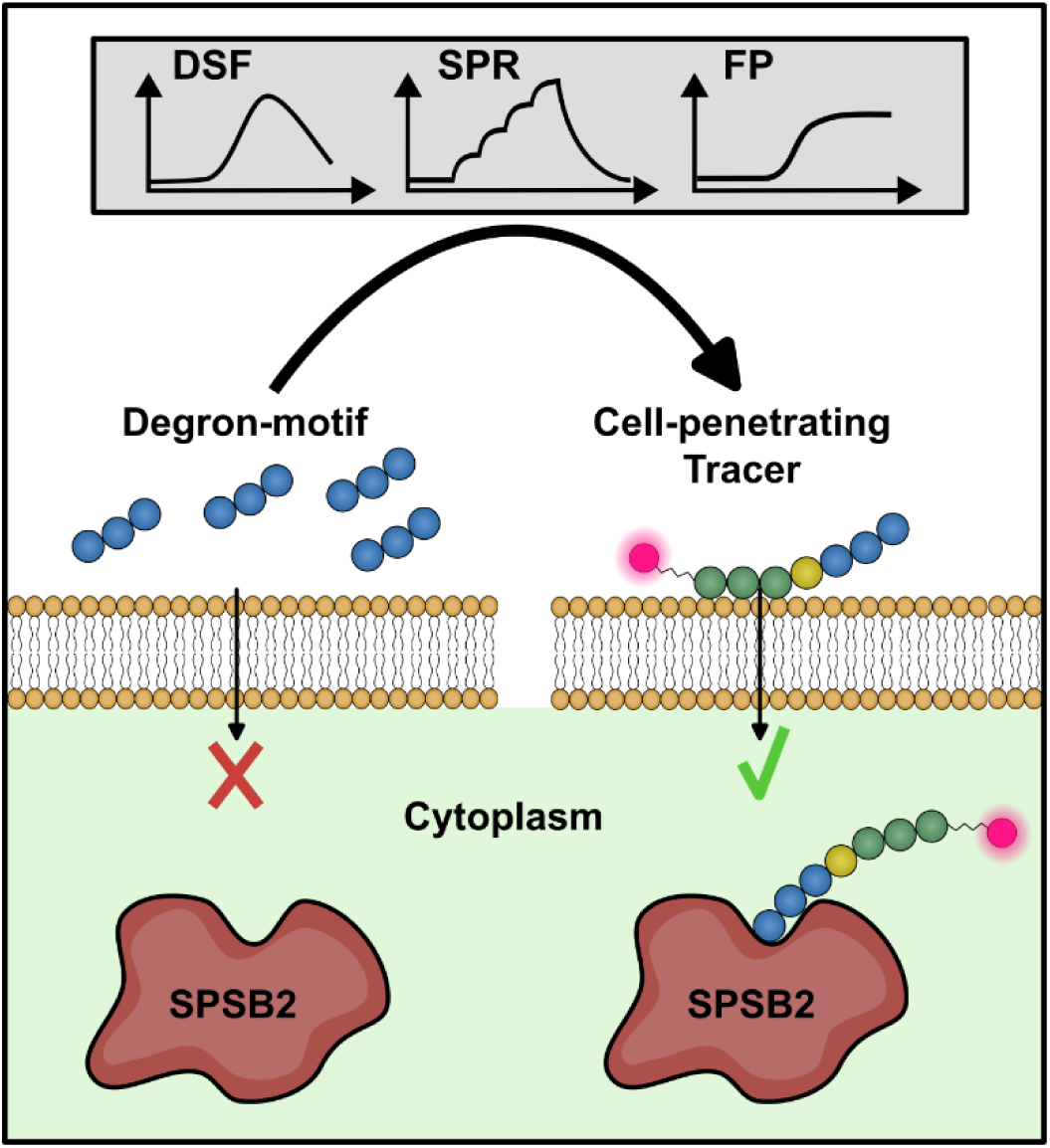

## Introduction

In pharmaceutical research, the implementation of various assay formats is critical for both *in vitro* and *in vivo* screening, as well as the characterization of potential ligands to target proteins. High-throughput *in vitro* assays which often utilize truncated target-binding domains, can lead to the discovery of high-affinity hits and potent lead structures. However, even with the most extensive screening efforts, most proteins are still considered non-druggable due to either not validated hit matter or failed optimization of lead compounds. [1, 2]

Peptidomimetics based on protein-protein interactions (PPIs) frequently serve as one of the most viable starting points for the development of inhibitors targeting these difficult-to-target domains. [3] E3 ligases fall into this category and new potent ligands for these domains would advance the emerging field of targeted protein degradation (TPD), a novel approach in drug discovery. In TPD, E3 ubiquitin ligases are recruited using heterobifunctional molecules such as PROteolysis TArgeting Chimeras (PROTACs), which bring the ligase into proximity with a target protein to induce polyubiquitination and subsequent proteasomal degradation. [4] PROTACs typically consist of a ligand (or “handle”) that binds the E3 ligase, linked to a warhead interacting with the target protein, thereby inducing proximity of the target with the ubiquitin degradation machinery. Thus, PROTACs hijack the natural degradation pathway, in which E3 ligases recognize short linear motifs known as degron sequences within proteins. [5, 6] These inherent degrons represent especially promising templates for designing effective PROTAC-handles. Recent examples on this topic have been published as studies on the discovery of first peptide-based compounds targeting the PRYSPRY domain of E3 ubiquitin ligase TRIM7 or the kelch domain of E3 ligase KLHDC2. [7, 8] Although peptide-based molecules show promise as lead structures, their often hydrophilic nature and polar properties limit membrane permeability and thus hinders studies on full-length proteins in cells. [9]

SPSB2 (SPRY domain-containing SOCS box protein 2) is a prime example for a challenging drug-target with a polar degron motif. SPSB2 acts as an adaptor protein in the Elongin B/C-Cullin-5-SPRY domain and SOCS box (ECS) E3 ubiquitin ligase complex, facilitating the ubiquitination and subsequent proteasomal degradation of target proteins harbouring the SPSB2 degron sequence. [10] For instance, degron-mediated interaction with inducible nitric oxide synthase (iNOS) and the SPRY domain (SPSB2^SPRY^), results in iNOS degradation, thereby modulating nitric oxide (NO) production in immune responses. Inhibiting the SPSB2-iNOS interaction may enhance iNOS activity, presenting a potential therapeutic strategy for chronic infections. [10] Beyond immune regulation, SPSB2 also exhibits antiviral activity by targeting the hepatitis-C-virus non-structural protein NS5A for ubiquitination and degradation, ultimately suppressing viral replication. [11] The involvement of SPSB2 in various essential cellular processes makes it an attractive target for drug development aimed at regulating protein degradation pathways. Given its role in protein ubiquitination, it is additionally considered a promising target for PROTAC development to achieve selective protein degradation. [12]

The available crystal structures and biophysical data of SPSB2^SPRY^ in complex with target peptides provide valuable insights into its substrate recognition and potential avenues for drug design. [12, 13] Yet to date, only potent binders incorporating the charged peptidic binding motif “DINNN”, like Peptide-Natural product-inspired hybrids (PepNats) or cyclized peptidomimetic ligands, have been reported. [14, 13, 15] The charged nature of this motif has posed significant challenges for cellular uptake. To address this, You et al. proposed a strategy to enhance internalization of these potent binders by conjugating an RGD motif to the peptide, thereby potentially facilitating integrin-dependent endocytosis. [14] This group also studied internalization by a cell-penetrating arginine-rich peptide (RRRRRRRRR) fused to the iNOS-based interaction sequence, ultimately inducing increased NO production. [16] Another recent study published in 2022 employed a similar strategy by conjugating a fluorophore (Cy5) and a cyclic cell-penetrating peptide to the iNOS binding sequence, enhancing NO production and enabling visualization of cellular uptake. [17]

In this study, we used SPSB2 as a representative PPI/E3 ligase target with a highly polar degron motif to establish functional *in vitro* assays that can be easily adapted to *in cellulo* target engagement formats through fluorophore conjugation. We introduce a versatile strategy to enable cellular engagement studies of otherwise non-permeable E3 degron peptide sequences by adding polycationic cell-penetrating peptides (CPPs) to facilitate efficient intracellular delivery. Using confocal microscopy, we evaluated various CPP motifs to assess their cellular uptake efficiency and localization. To determine the specific interaction between the modified degron peptide and the full-length SPSB2 protein, we conducted nano bioluminescence energy transfer (NanoBRET) experiments, developing a qualitative displacement assay for interaction studies in cellular environments. Furthermore, we assessed the cytotoxicity by monitoring cell viability, excluding potential aggregation-related effects of the CPP-conjugated peptides and fluorophore-linked tracer molecules. Here, we present a study that offers a simple, yet effective approach to study and characterize non-permeable degron motifs of E3 ligases using SPSB2 as a representative example.

## Results and Discussion

### Screening of SPSB2^SPRY^-binding Ligands

In an effort to identify new binding scaffolds for SPSB2, we subjected the C-terminal SPRY-domain of the protein to various high-throughput screening assays, including E-ASMS (>8,200 compounds) and virtual docking of an in-house library (>7,500 compounds). [18] Initial assessment of ligand binding was carried out with SPSB2^SPRY^ using Surface Plasmon Resonance (SPR). For additional biophysical characterization, thermal shift assays, also known as Differential Scanning Fluorimetry (DSF) and Nuclear Magnetic Resonance (NMR) were performed. Unfortunately, our biophysical evaluation of the potential screening hits did not confirm our hit matter, despite the existence of various published peptides with nanomolar affinity for SPSB2^SPRY^. [19, 20] Therefore, the focus of our study was the establishment of a cellular binding/competition assay for SPSB2.

### Strategy for the Design and Evaluation of Cell Penetrating Peptides

Biophysical *in vitro* assays are essential for ligand assessment, as they can be used to determine binding affinity and characterize structural interactions with the target protein. However, they do not assess membrane permeability and are rarely applied to full-length proteins, both of which are critical factors influencing ligand evaluation. To address these limitations in screening campaigns, we chose human SPSB2 (hSPSB2) as a representative E3 ligase binding a highly polar non-permeable C-terminal peptide degron. We used various biophysical in vitro assays for initial peptide/tracer selection and subsequently developed a cellular screening approach by combining two techniques: I) confocal imaging to monitor peptide/tracer permeation into the cell and II) NanoBRET to assess interactions with the full-length target protein SPSB2. The synthesized peptides were designed with an N-terminal CPP motif followed by a C-terminal DINNN peptide binding sequence. For the generation of fluorescent tracer molecules, BODIPY 576/589 was additionally conjugated to the N-terminus, as this dye was required for establishing NanoBRET as an in vivo target engagement assay. (Figure 1A) To compare different CPPs, we investigated the use of an RGD motif, which is known to be internalized via endocytosis as reported by You et al. (2017), and the TAT motif (YGRKKRRQRRR), a well-characterized and frequently used cell-penetrating peptide. [21, 22] For the TAT construct, we additionally designed a cyclized variant by adding a disulfide bridge flanking the TAT sequence to improve cell permeability. [22] Linear peptides are typically unstructured in solution and susceptible to degradation by both endo- and exopeptidases due to their free N- and C-termini. In contrast, cyclic peptides offer greater stability against enzymatic degradation and often exhibit enhanced binding affinity and in vivo potency, making them more attractive for assay development. [23] However, the design and synthesis of cyclic peptides can be technically challenging and costly, which is why linear peptides remain predominant in most CPP-related biological studies.

**Figure 1.**
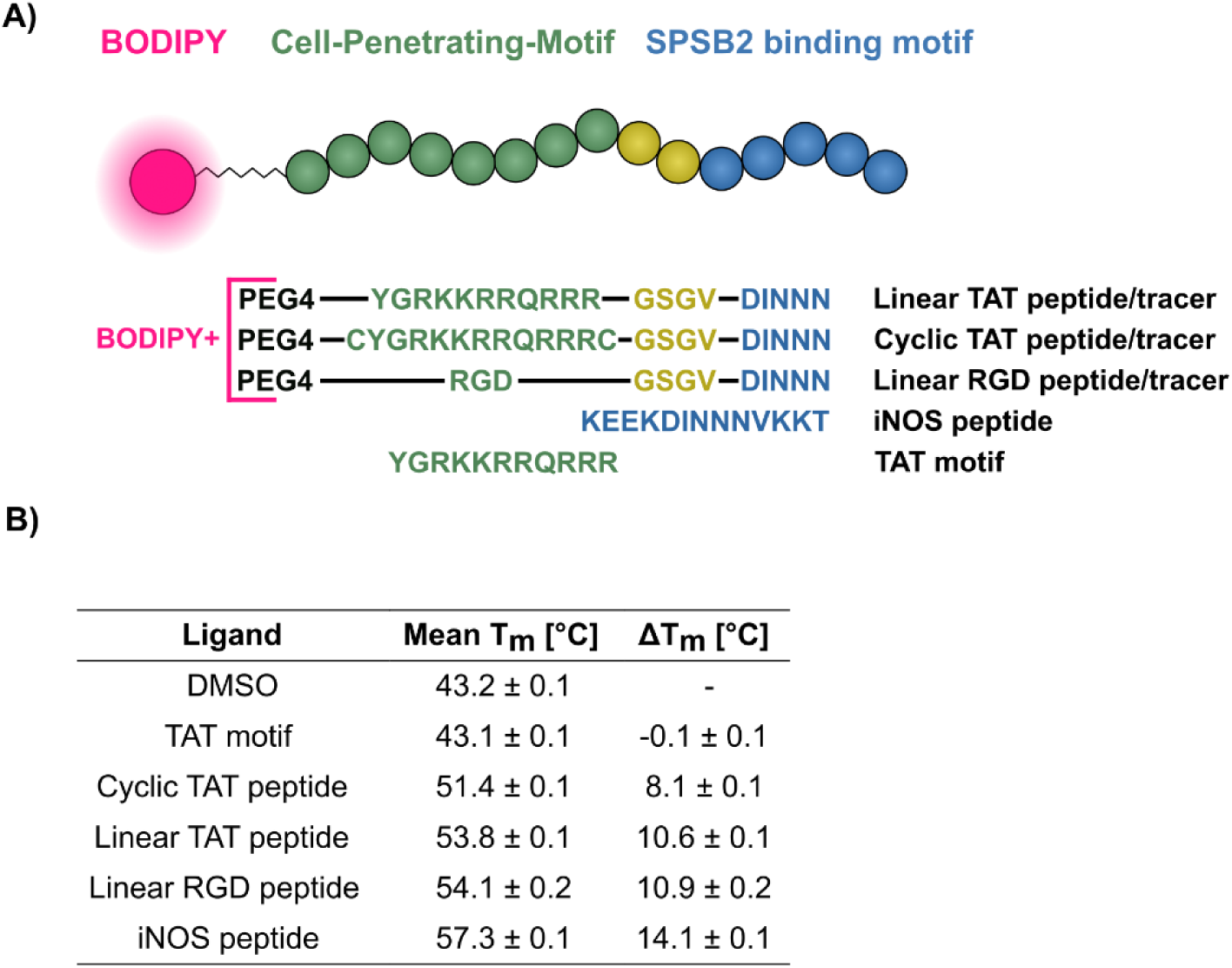
Peptide/tracer design strategy and overview of thermal shift analyses results. (A) Overview of the designed peptides and tracer constructs. From top to bottom: linear TAT peptide/tracer, cyclic TAT peptide/tracer, linear RGD peptide/tracer, iNOS peptide, and TAT motif. (B) DSF results of the selected peptides against hSPSB2^SPRY^, Tm and ΔTm values are depicted as mean ± SD (n = 4).

### Degron Sequences are Potent Ligands of the SPSB2 SPRY Domain

To investigate possible interference of the flanking cell penetration tags on DINNN binding to hSPSB2^SPRY^, we first carried out thermal shift analyses, which revealed significant differences in melting temperature between the different peptides (Figure 1B). As expected, the native iNOS interaction motif KEEKDINNNVKKT (iNOS peptide) exhibited a strong ΔTm of 14.1 ± 0.1 °C in agreement with the published nanomolar affinity of this degron. [13] The N-terminal presence of CPP sequences conjugated to the DINNN motif displayed a decrease in ΔTm to 10.9 ± 0.2 °C and 10.6 ± 0.1 °C for the linear RGD peptide (PEG4-RGDGSGVDINNN) and linear TAT peptide (PEG4-YGRKKRRQRRRGSGVDINNN), respectively. The ΔTm for the cyclic TAT peptide variant (PEG4-CYGRKKRRQRRRCGSGVDINNN) also decreased to 8.1 ± 0.1 °C. As a control experiment, we demonstrated that the isolated TAT motif did not affect the melting temperature of hSPSB2^SPRY^. These results imply prominent binding affinity differences among the peptides induced by the presence of different CPP motifs. Therefore, we evaluated the DSF binding results via orthogonal biophysical techniques using single cycle kinetics analysis in SPR. (Figure 2A, Supplement Table 1).

**Figure 2.**
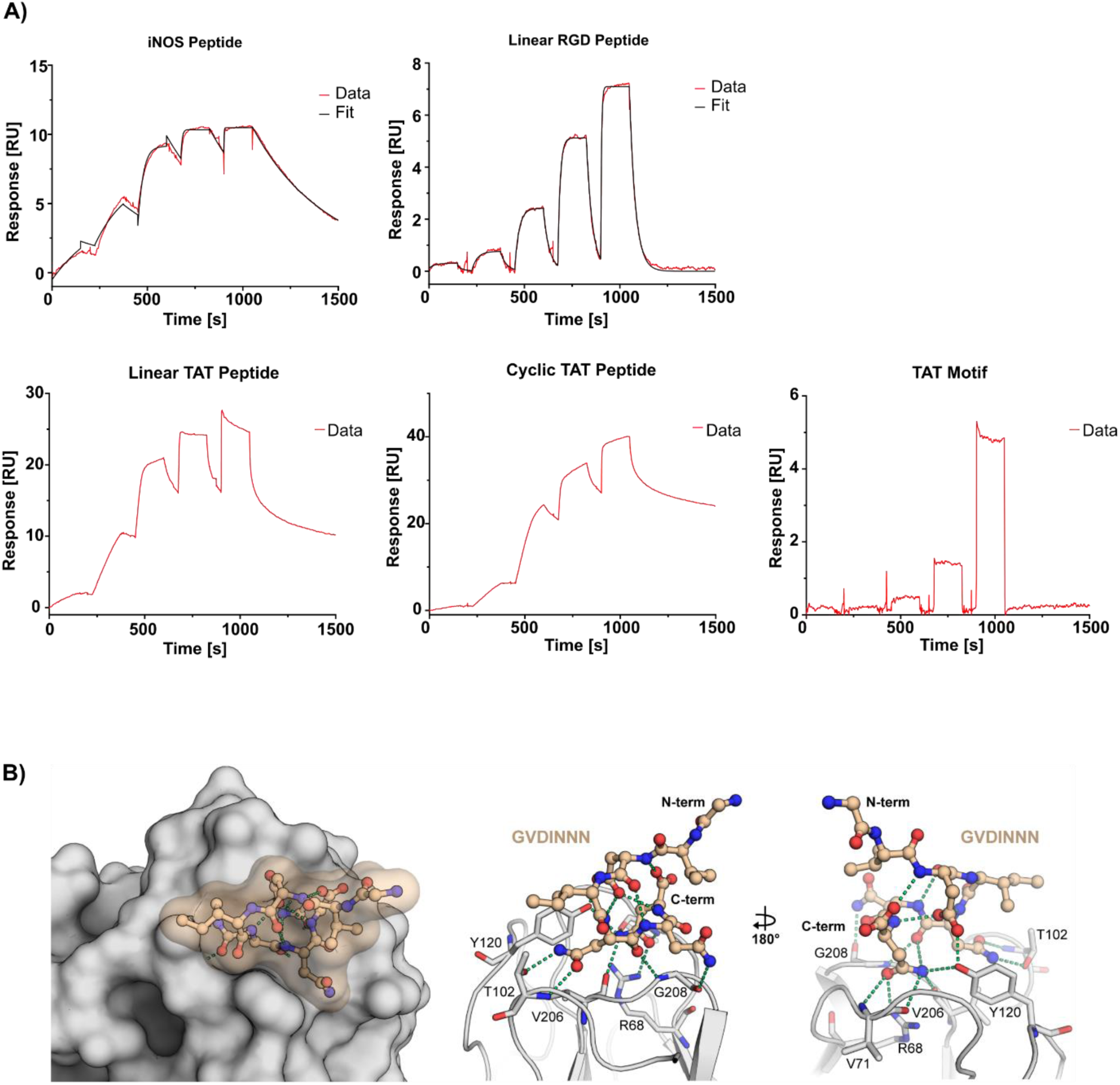
Biophysical SPR evaluation and structure of hSPSB2^SPRY^ in complex with linear TAT peptide. (A) Surface plasmon resonance single cycle kinetics sensorgrams of iNOS peptide, linear RGD peptide, linear TAT peptide, cyclic TAT peptide and TAT motif. (B) Crystal structure of hSPSB2^SPRY^ co-crystallized with linear TAT peptide (PDB: 9RV5). Hydrogen bonds and polar interactions of hSPSB2^SPRY^ with GVDINNN are shown as green dotted lines.

As expected, all peptides showed high affinity binding to immobilized SPRY domain of SPSB2. The published iNOS peptide exhibited a K_D_ of 0.8 ± 0.1 nM, while the dissociation rate constant (k_off_) could be determined as (2.4 ± 0.1)*10^-3^ s^-1^ and the association rate constant (k_on_) as (2.9 ± 0.1)*10^6^ M^-1^s^-1^. In comparison, the peptide containing the RGD motif displayed a reduced affinity of 162.0 ± 4.0 nM and more than 10x faster kinetics with a k_off_ of (3.2 ± 0.1)*10^-2^ s^-1^ and a k_on_ of (2.0 ± 0.1)*10^5^ M^-1^s^-1^ which was in agreement with our previous DSF results. Intriguingly, while the TAT motif itself did not seem to interact specifically or only very weakly with hSPSB2^SPRY^, the linear as well as cyclic TAT peptide exhibited a non 1:1 binding interaction as implied by an apparent two-phase binding behaviour, characterized by a slow dissociation phase following an initial fast dissociation phase.

Visual assessment of the data suggests a much longer residence time for both TAT peptides as compared to the linear RGD and iNOS peptide. However, the complexity of a non 1:1 binding interaction complicates direct quantitative comparison with systems exhibiting a 1:1 binding behaviour. We therefore relied solely on a qualitative comparison rather than determining artificial kinetic or affinity values for each TAT peptide. A comparable discovery was reported by Rahman et al. in 2022 for murine SPSB2^SPRY^, who used a similar CPP-analogue conjugated to the interaction motif KDINNNV. Although they were also unable to determine a K_D_ value for the peptide, they observed a significantly reduced SPR off-rate, possibly resulting from additional interactions with the cationic residues of the CPP-moiety. [17]

### High Affinity Degrons Share Similar Binding Modes

To further investigate the influence of the different CPP peptides towards the DINNN-hSPSB2^SPRY^ interaction at atomic level, we applied Crystallography (X-ray) and NMR. Fortunately, we were able to determine a high-resolution (1.75 Å) co-crystal structure of the linear TAT peptide in complex with hSPSB2^SPRY^ (PDB ID: 9RV5, Figure 2B, left panel). Similar to previous reports, we did not observe interpretable electron density for the entire peptide but obtained only a well-defined density for the C-terminal sequence GVDINNN. Consistent with earlier findings, the recognition sequence binds to the SPRY domain identically, as evident from its superposition with PDB entry 6KEY (Figure S1). The degron peptide formed polar interactions with the backbone amides of V71, T102, V206, and G208, as well as with the side chains of R68, T102, and Y120 (Figure 2B, right panel). Furthermore, the peptide conformation was stabilized by an internal network of hydrogen bonds (Figure 2B, right panel). The absence of an electron density for the TAT sequence in our structure suggested a rather flexible and dynamic or transient binding.

To further visualize potential differences in binding, we tested the iNOS peptide as well as the linear TAT peptide against ^15^N-labeled hSPSB2^SPRY^ and conducted 2D ^1^H-^15^N correlation NMR experiments. While both peptides induced substantial chemical shift perturbations relative to the apo protein spectrum, several resonances showed different chemical shift changes upon binding of the linear TAT peptide compared to the iNOS-derived peptide. (Figure S2) These data suggested distinct interaction patterns of the backbone amides with each peptide. The spectral differences aligned with our SPR data, where the TAT peptide demonstrated altered dissociation kinetics and a non-1:1 binding behaviour, consistent with potentially different conformational dynamics upon binding. Nonetheless, due to the lack of an NMR assignment for hSPSB2^SPRY^, it was not possible to determine which exact residues were affected by peptide binding.

The linear RGD peptide, linear TAT peptide and cyclic TAT peptide were subsequently used to develop tracer molecules (Table 1) for initial fluorescence polarization (FP) assays as an additional system to compare hSPSB2^SPRY^ binding affinities (Figure 3). As previously described, fluorescent tracers were designed by conjugating BODIPY 576/589 to the N-terminus of each CPP-conjugated peptide and were evaluated via FP target engagement by titrating 80 pM - 10 µM hSPSB2^SPRY^ to 4 nM tracer. (Table 1) As expected, each tracer exhibited nanomolar affinities to hSPSB2^SPRY^, while the linear RGD tracer displayed the weakest affinity of ∼56 nM in line with our previous SPR results. Interestingly, while the affinity of the cyclic TAT tracer was determined as ∼0.6 nM, the K_D_ of the linear TAT tracer was higher at ∼13 nM. While these results did not fully align with our DSF results, the substantially higher affinity of the cyclic TAT was consistent with our expectations, as cyclic peptides generally offer enhanced binding affinities due to their reduced conformational flexibility and structural organization compared to their linear counterparts [23].

**Figure 3.**
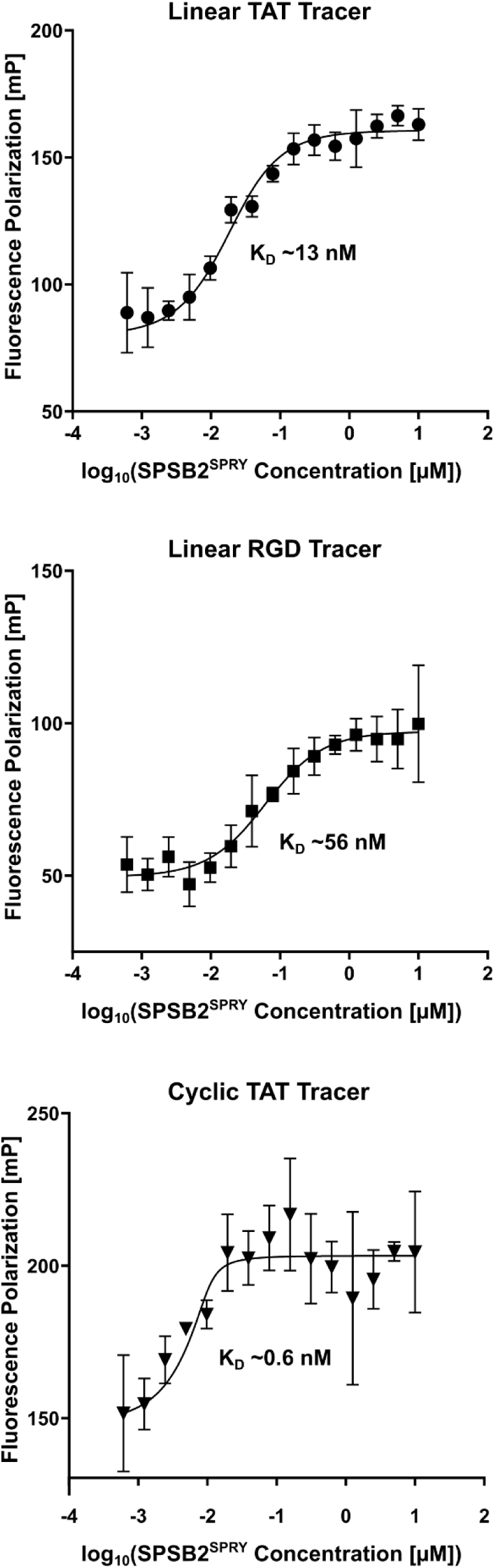
*In vitro* target engagement of tracer molecules. Fluorescence polarization tracer titration of linear TAT tracer, linear RGD tracer and cyclic TAT tracer with respective KD values. Measured values are depicted as mean ± SD (n = 3).

**Table 1.**
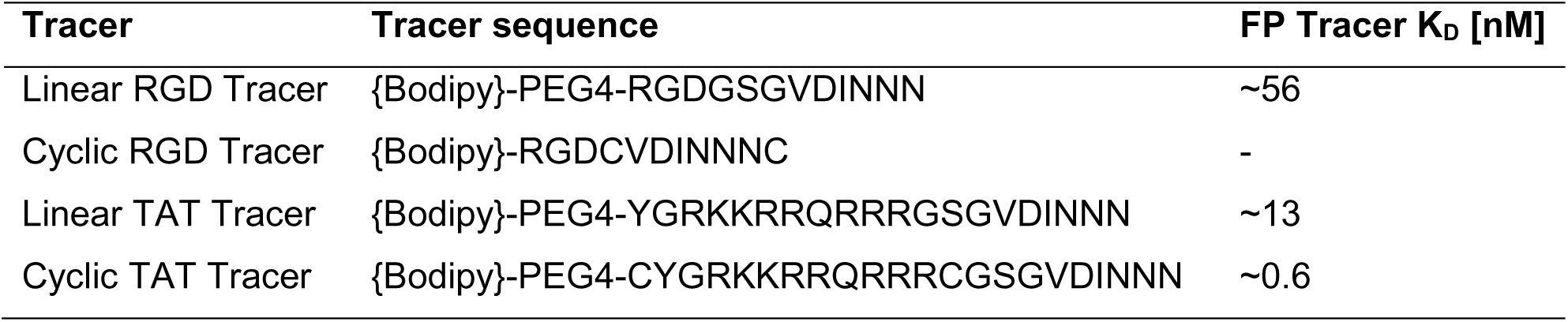
Overview of synthesized tracer molecules linear RGD tracer, cyclic RGD tracer, linear TAT tracer and cyclic TAT tracer.

| Tracer | Tracer sequence | FP Tracer $K_D$ [nM] |
| --- | --- | --- |
| Linear RGD Tracer | {Bodipy}-PEG4-RGDGSGVDINNN | ~56 |
| Cyclic RGD Tracer | {Bodipy}-RGDCVDINNNC | - |
| Linear TAT Tracer | {Bodipy}-PEG4-YGRKKRRQRRRGSGVDINNN | ~13 |
| Cyclic TAT Tracer | {Bodipy}-PEG4-CYGRKKRRQRRRCGSGVDINNN | ~0.6 |

### Discovery of Cell-permeable Tracer Molecules

To check the permeability of each tracer, we tested them at varying concentrations from 400 nM to 50 µM in colon cancer cells (HCT116) and assessed the fluorescent signal at four different time points from 30 min to 210 min. We additionally tested a fourth short tracer incorporating the RGD motif as a cyclic RGD tracer (Table 1). Cyclization of this tracer was achieved through disulfide bridges flanking the protein interaction sequence to increase stability against enzymatic degradation and enhance potential binding affinity to SPSB2. [23]

No visible uptake of our RGD-conjugated tracers was detected, even though HCT116 cells, as colon cancer cells, express high levels of integrins and RGD-based motifs have been proposed as a viable strategy for cell permeation. [24, 14] This highlights the complexity and challenges associated with designing functional, cell-permeable molecules, as not every well-characterized motif reliably facilitates cellular permeation in different experimental settings. In contrast, both TAT-conjugated tracers demonstrated visible cellular permeability and cellular localization at tracer concentrations of around 1 µM (Figure 4A). Accumulation of both tracers at the cellular membrane was evident after 30 min, and fluorescence corresponding to the peptide dye conjugate became clearly visible in the cellular cytoplasm after 90 min of incubation time. The fluorescence intensity in the cytoplasm continued to increase over time, while displaying changes in the cell membrane morphology, potentially resulting from TAT-based crowding effects, endocytosis and transient pore formation. [25–30] Given that the cyclic form of the TAT sequence exhibited increased cellular permeability, and that our FP study also demonstrated higher target affinity compared to its linear counterpart, we selected the cyclic TAT variant for subsequent assay development [22, 31]. To exclude the possibility of cell-membrane disruption or other cytotoxicity inducing effects potentially caused by the peptide or dye-associated tracer, we tested their cytotoxicity at concentrations ranging from 0.39 µM to 50 µM at 2 h incubation, using a CellTiter-Glo assay (Figure 4B). The cyclic TAT peptide had no impact on cell viability at any tested concentration, suggesting that cellular functions and the plasma membrane remained intact. In contrast, the corresponding tracer led to reduced cell survival at concentrations above 10 µM, potentially due to solubility limitations and aggregation effects. These results suggested that using tracer concentrations well below 10 µM is unlikely to disrupt cellular integrity and can therefore be readily applied to cellular target engagement assays.

**Figure 4.**
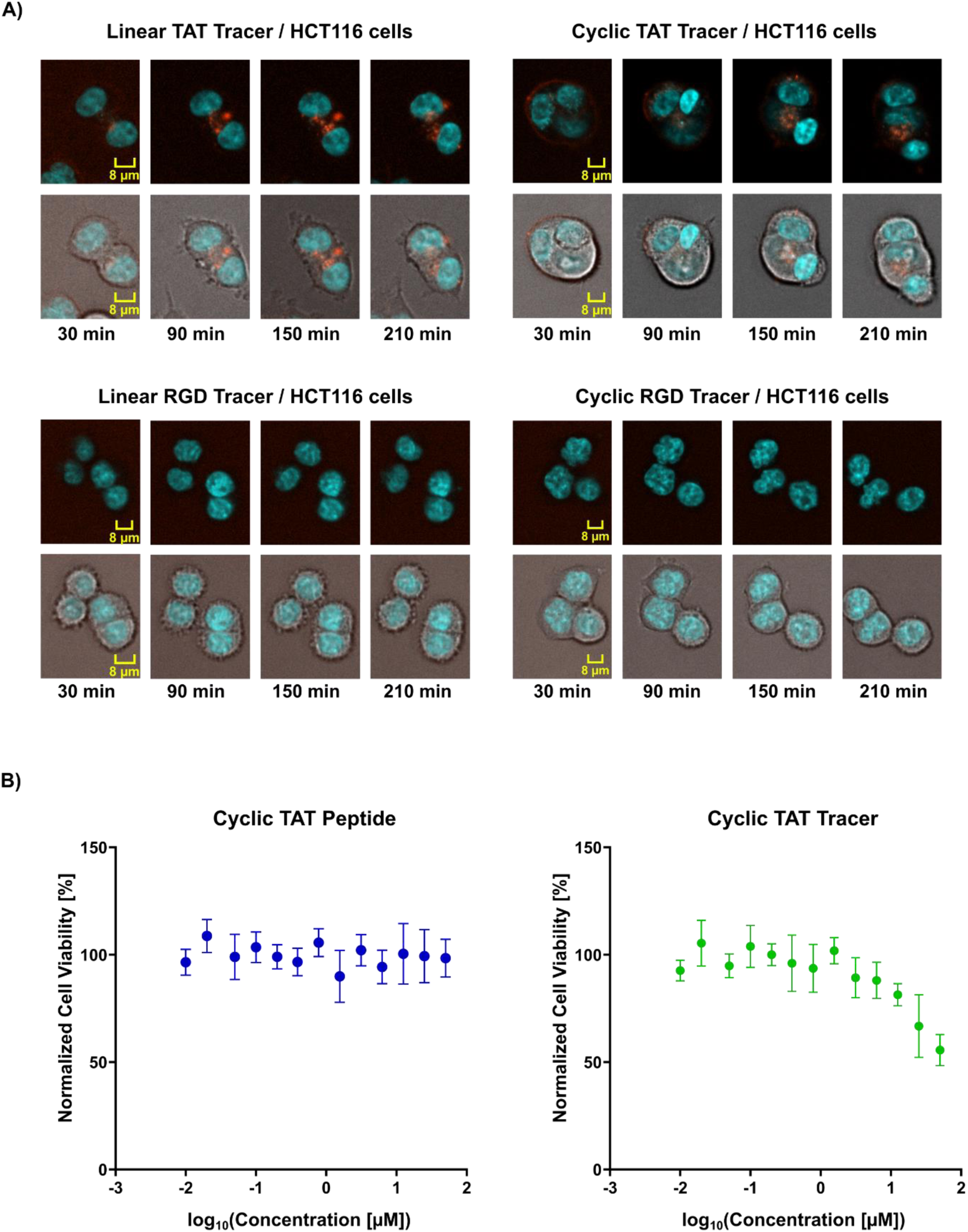
Cellular permeability and cytotoxicity studies. (A) Confocal imaging of HCT116 cells incubated with 1.56 µM tracer and imaged at 30 min, 90 min, 150 min and 210 min. Images were acquired using brightfield and blue/red fluorescence channels. Tracer permeation is visualized in red: linear TAT tracer (top left), cyclic TAT tracer (top right), linear RGD tracer (bottom left), and cyclic RGD tracer (bottom right). Nuclei were stained with Hoechst dye (blue). (B) CellTiter-Glo analysis of cyclic TAT peptide (left) and cyclic TAT tracer (right). Measured values are depicted as mean ± SD (n = 6).

### Cellular Assay Development by Degron-Peptide Displacement

We further conducted NanoBRET assays, by titrating cyclic TAT tracer in a dose-dependent manner and assessing target engagement in both intact cells (Figure 5A, left) as well as digitonin-permeabilized cells (Figure 5A, right). The K_Dapp_ value of the tracer was determined as 86 nM for intact cells and 346 nM for permeabilized cells. For both conditions, no signal saturation was observed at the highest tested tracer concentrations. However, this lack of saturation has been reported many times and it may only reflect the approaching of solubility limits (https://www.tracerdb.org/). [32] Importantly, many of these assays were performing well, as long as a measurable BRET signal was maintained and displacement with unlabelled compounds was observed. Gratifyingly, dose-dependent titration of the competing cyclic TAT peptide demonstrated clear displacement in permeabilized (Figure 5B, right) as well as intact cells (Figure 5B, left). However, a high signal variability for intact cells led to inconsistencies in defining a clear intact cell EC_50_. To address this, we repeated the assay as biological replicates at 20 nM and 30 nM tracer concentrations, confirming displacement reproducibility (Supplement Figure S3).

**Figure 5.**
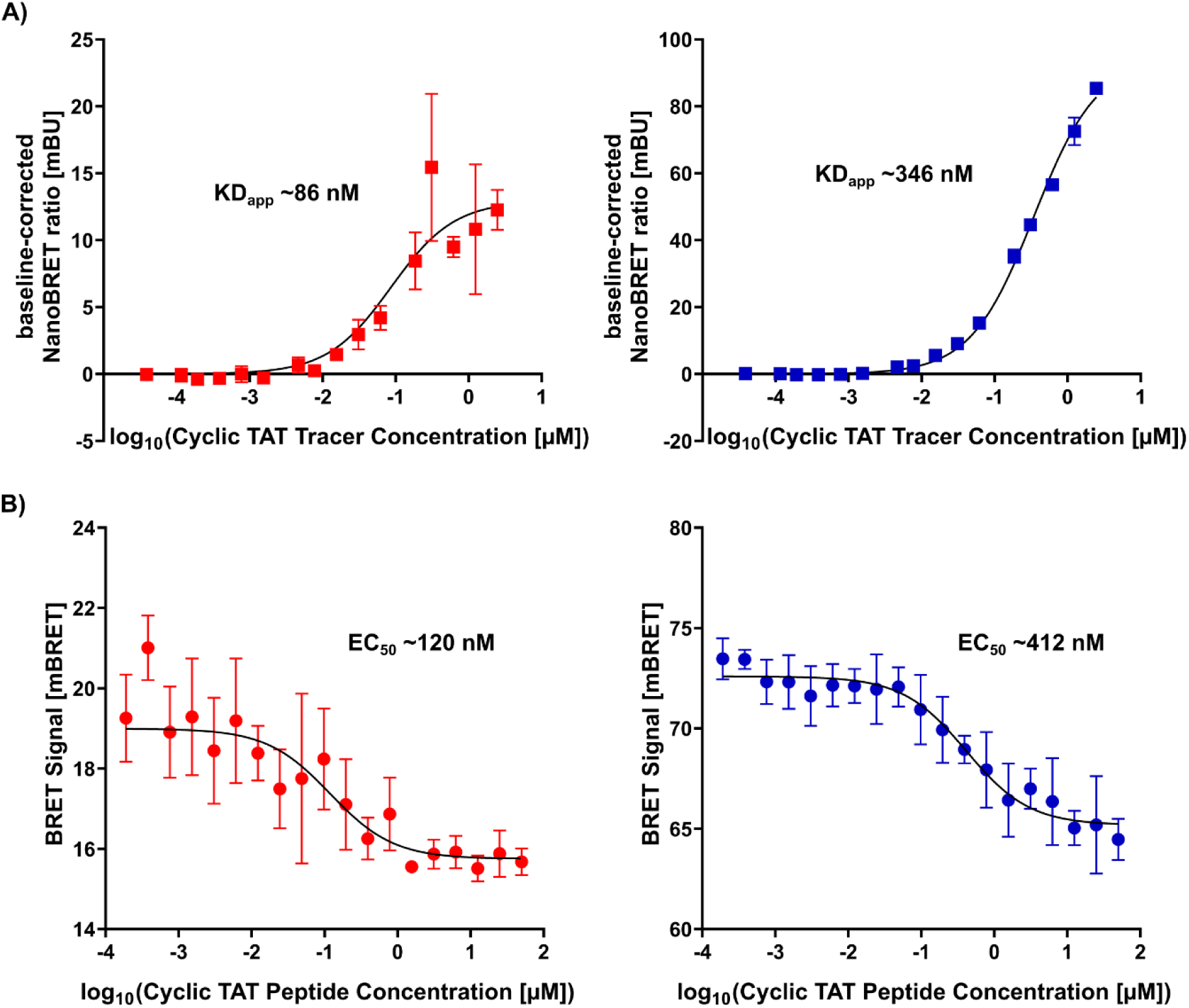
Cellular target engagement and displacement of fluorophore-tagged cyclic TAT tracer. (A) NanoBRET tracer titration of cyclic TAT tracer in intact HCT116 cells (left) and permeabilized cells (right) with KDapp values determined for both conditions. (B) Displacement of cyclic TAT tracer with cyclic TAT peptide in intact (left) and permeabilized cells (right) and their respective EC50 values. Measured values are shown as mean ± SD (n = 3).

As the signal variability increased with higher tracer concentrations in intact cells, careful selection of appropriate tracer concentrations was necessary to maintain signal strength while minimizing deviations. This effect is likely attributed to the complex uptake mechanism mediated by the TAT sequence, as both the tracer and the competing peptide rely on the same internalization pathway. Additionally, while the TAT-based tracer was efficiently taken up by HCT116 cells, as shown by confocal microscopy, previous studies suggested that escape across endosomal membranes and subsequent cytosolic delivery of TAT-conjugates can be very inefficient, potentially limiting the amount available for effective target engagement and leading to an inhomogeneous tracer distribution. [30] While careful evaluation of the results was necessary, clear displacement of the tracer was observed in cells, enabling at least qualitative screening of ligands in intact cells as well as quantitative affinity determination using the full-length protein in permeabilized cells.

## Outlook

In this study, we demonstrate the use of cell-penetrating peptides for ligand discovery, targeting the E3 ligase SPSB2 with a polar non-permeable degron peptide binding motif. The combination of two techniques I) confocal imaging and II) NanoBRET enabled the development of a qualitative cellular screening approach for SPSB2. Moreover, biophysical characterization of the synthesized peptides with CPPs and the dye motif elucidated the influence of these modifications on the interaction with the target protein. The peptides and their binding modes may serve as templates for the design of peptidomimetic ligands for SPSB2, as already demonstrated for TRIM7 by Muñoz Sosa and Lenz et al. in 2024. [7]

The strategies and insights gained from this work may be broadly applicable to other PPI targets beyond E3 ligases, thus facilitating cellular screening of proteins that are otherwise difficult to engage. Additionally, the introduced CPP-fluorophore conjugation approach may support pilot studies of poorly characterized E3 ligases with understudied degron motifs, or even help in the cellular delivery of otherwise impermeable small-molecule PROTACs for early-stage TPD studies. We hope the results presented in this study will help leverage research aimed at developing effective, cell-permeable molecules to engage targets that are currently considered undruggable, thereby advancing their potential ligandability.

## Methods

### Protein Expression and Purification

Human His_6_-SPSB2^SPRY^ (86-219) was expressed as a fusion construct in the *Escherichia coli* BL21(DE3) strain. Protein expression was performed by inducing cells cultured in LB-medium with 1 mM Isopropyl-β-D-thiogalactopyranosid (IPTG) at an OD_600_ = 0.6-1.0, followed by incubation overnight at 18 °C. Cell lysate was purified by affinity chromatography using HisTrap HP columns (Cytiva), followed by further purification via size-exclusion chromatography using a HiLoad™ 26/600 Superdex™ 75 pg column (Cytiva). The protein was finally stored in 30 mM HEPES pH 7.5, 100 mM NaCl and 0.5 mM TCEP.

### Peptide and Tracer Synthesis

Peptides and Tracer molecules were custom synthesized by Synpel Chemical, GeneCust and GenScript Biotech.

### Differential Scanning Fluorimetry

For peptide screening, 50 µM peptide was added to 5 µM His-hSPSB2^SPRY^ diluted in buffer containing 30 mM HEPES pH 7.5, 100 mM NaCl, 1 mM TCEP, 5x SYPRO Orange and 5% DMSO as quadruplicates. Thermal shift analyses were carried out using a QuantStudio 5 Real-Time PCR System (Applied Biosystems) with a temperature gradient ranging from 25 °C to 95 °C at an increase of 0.05 °C/s. Data analysis was performed by fitting a Boltzmann sigmoidal curve to the raw values using the Protein Thermal Shift software (Applied Biosystems).

### Crystallography, Data Collection and Refinement

Purified human SPSB2 SPRY domain at 3.5 mg/ml in SEC buffer was mixed with linear TAT-peptide (10 mM stock solution in DMSO) to a final concentration of approximately 500 µM (final DMSO conc. 5%). This protein-peptide complex was co-crystallized at 20 °C using the sitting-drop vapour diffusion method in a 1:1 ratio with a reservoir solution containing 22% PEG3350, 10% ethylene glycol and 0.2 M sodium nitrate. Crystals of hSPSB2^SPRY^ grew to full size within 1-3 days and were later determined to belong to space group P2_1_, with two SPSB2-peptide complexes per asymmetric unit (AU). Before flash-freezing the crystals, the ethylene glycol concentration was raised to 25% for cryo-protection.

Diffraction data were collected at Diamond Light Source I03 (Didcot, UK) at a wavelength of 0.97625 Å at 100 K. Data were automatically processed using xia2 [33] and scaled with aimless [34]. The PDB structure with the accession code 3EMW [19] was used as an initial search model for molecular replacement using the program MOLREP [35]. The final model was built manually using Coot [36] and refined with REFMAC5 [37], which is a part of the CCP4 suite [38]. Data collection and refinement statistics are summarized in Supplement Table 2.

### Surface Plasmon Resonance

Surface Plasmon Resonance experiments were conducted on a Biacore T200 instrument with a Series S Sensor Chip CM5 (Cytiva) and 10 mM HEPES pH 7.5, 150 mM NaCl, 0.5 mM TCEP, 0.05% Tween20 and 2% DMSO for the running buffer. His-hSPSB2^SPRY^ was diluted to 5 µg/mL in Acetate pH 5.5 and immobilized at 10 µL/min to reach 400-700 RU response levels on Flow-channel (FC) 2-4 for triplicates. Immobilization was performed via standard amine coupling procedure by injecting a mixture of 483 mM EDC and 10 mM NHS for activation with a subsequent injection of 1 M ethanolamine for deactivation. FC1 was used as a reference surface without protein-injection. Kinetic titration experiments were done using single cycle kinetics procedure by injecting 5 concentrations of peptide at 30 µL/min ranging from 0.8 nM to 1 µM. Following each peptide titration, multiple buffer injections were performed to reach a stable baseline. Solvent correction and blank injections were also included. After referencing, blank subtraction, and solvent correction, the data were analyzed using the Biacore T200 evaluation software. Sensorgrams with overlayed fit were replotted using GraphPad Prism version 8.0.1.

### Fluorescence Polarization Assay

Fluorescence Polarization experiments were performed as triplicates in FP buffer (50 mM HEPES pH 7.5, 250 mM NaCl, 2 mM TCEP, 0.05% Tween-20 and 2% DMSO). For tracer titration assays, protein concentrations ranging from 80 pM to 10 µM were titrated to 4 nM tracer in a 384-well flat-bottom plate (Greiner Bio-One, #784076), incubated for 40 minutes, and measured using a PHERAstar FSX plate reader equipped with a Transcreener-specific FP optic module (FP 590/675/675). K_D_ values [µM] for each tracer were determined by fitting the data using a non-linear regression model accounting for ligand depletion, via GraphPad Prism version 8.0.1.:

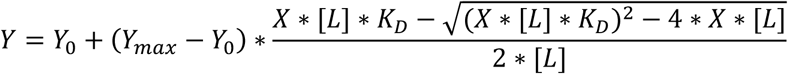

Here, Y is the measured fluorescence intensity [mP], X is the protein concentration [µM], Y_0_ is the signal intensity of the free tracer [mP], Ymax is the signal intensity of the fully bound tracer [mP] and [L] is the tracer concentration [µM].

### NanoBRET Interaction Assay

NanoBRET experiments were conducted by transfecting HCT116 cells 24 hours before the measurement with C-terminally Nanoluc-tagged human full length SPSB2 using FuGENE HD Transfection Reagent (Promega). For tracer titration assays, 2.0*10^5^ cells in Opti-MEM I Reduced Serum Medium (Opti-MEM) were seeded into a white 384-well flat-bottom plate (Greiner Bio-One, #784075). Afterwards, 39 pM - 2.5 µM tracer was titrated to the transfected cells as triplicates at 1% DMSO using an ECHO 550 acoustic dispenser, then incubated for 2 h at 37 °C and 5% CO_2_. For displacement assays, 20 nM and 30 nM of cyclic TAT tracer as well as varying concentrations of peptide ranging from 6 nM to 50 µM were titrated to the cells as triplicates using an ECHO 550 acoustic dispenser. To permeabilize cells, 25 µM digitonin was added to each well. BRET signal was measured on a PheraSTAR FSX plate reader with a luminescence filter pair (450 nm BP filter and 610 nm LP filter). Data evaluation was carried out with GraphPad Prism version 8.0.1. For tracer titration, data was baseline-corrected, then fitted with the following non-linear equation to determine the K_Dapp_ of the tracer:

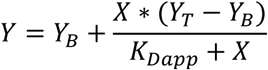

where Y is the baseline-corrected BRET signal [mBU], Y_B_ and Y_T_ is the baseline-corrected BRET signal at the lower and higher plateau, respectively and X corresponds to the tracer concentration [µM].

Displacement data was evaluated by fitting the BRET signal with a non-linear model to determine the EC_50_ of the displacing peptide:

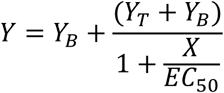

here, Y is the BRET signal [mBU], X is the concentration of the displacing peptide [µM], Y_B_ is the BRET signal [mBU] at the bottom plateau and Y_T_ corresponds to the BRET signal [mBU] at the top plateau.

### CQ1 Confocal Microscopy Permeation Study

HCT116 cells (ATCC: CCL-247) were cultured in DMEM plus L-glutamine (high glucose) supplemented by 10% FBS (Gibco) and 1% penicillin/streptomycin (Gibco). Cells were seeded at a density of 2000 cells per well in a 384-well plate (cell culture microplate, PS, f-bottom, μClear, 781091, Greiner) with a final volume of 40 µL. Cells were stained with 60 nM Hoechst 33342 (Thermo Scientific) and incubated overnight at 37 °C and 5% CO_2_ to allow for plate attachment. The following day, media was aspirated, cells were washed once with 1X Dulbecco’s Phosphate Buffered Saline (DPBS) and 40 µL of Opti-MEM was added to each well to minimise background fluorescence. Linear TAT tracer, cyclic TAT tracer, linear RGD tracer and cyclic RGD tracer were added at concentrations ranging from 0.39 µM to 50 µM at 1% DMSO using an ECHO 550 acoustic dispenser (Labcyte). Fluorescent activity of each tracer was recorded at 30 min, 90 min, 150 min and 210 min while incubating the cells at 37 °C and 5% CO_2_. Images were acquired using a CQ1 high-content confocal microscope (Yokogawa, Musashino, Japan) with the following setup: Ex 405 nm/Em 447/60 nm (500 ms, 50%) for Hoechst 33342, Ex 561 nm/Em 617/73 nm (100 ms, 40%) for the Tracer series and brightfield (50 ms, 80%). Data analysis was performed using CellPath-finder software (Yokogawa).

### Cytotoxicity assay

For cytotoxicity measurements, 10 µL HCT116 cells in Dulbecco’s Modified Eagle Medium (DMEM) were seeded at 2*10^5^ cells/mL into a white 384-well-flat-bottom plate. 10 nM to 50 µM tracer or peptide as well as 25 µM digitonin was added to the cells as hexaplicates at 1% DMSO via an ECHO 550 acoustic dispenser, then incubated for 2 h at 37 °C and 5% CO_2_. 10 µL CellTiter-Glo® 2.0 Reagent (Promega) was added to each well, followed by an incubation of 10 min at room temperature. Luminescence was measured, using a PheraSTAR FSX plate reader. Data was normalized to cells treated with 1% DMSO (100% survival control) and digitonin (0% survival control), then plotted using GraphPad Prism version 8.0.1.

### Nuclear Magnetic Resonance measurements

Two-dimensional ^1^H-^15^N correlation spectra were recorded on a Bruker Avance NEO 600 MHz spectrometer using a BEST-TROSY pulse sequence. [39] All measurement were performed at a sample temperature of 298 K and employed a cryogenic 5 mm ^1^H{^13^C/^15^N} triple-resonance (TCI) probe. Samples consisted of 62 µM ^15^N labelled hSPSB2^SPRY^ in 30 mM HEPES buffer (pH = 7.5) containing 100 mM NaCl, 1 mM TCEP, 0.15 mM DSS (as internal standard) and 5% D_2_O. Spectra were recorded of the free protein and of the complexes with either linear TAT peptide or iNOS peptide at a slight excess (82 µM). Acquisition times were 62.4 ms and 78.9 ms in the ^1^H and ^15^N dimensions, respectively. For each of 384 FIDs 32 transients were accumulated, resulting in measurement times of 1.5 h per spectrum, using a recycle delay of 0.3 s.

### Virtual Docking

Our kinase-focused in-house compound library of approximately 7,500 compounds was virtually screened against hSPSB2 (PDB ID: 6DN6) using AutoDock-GPU with default parameters. [40] The docking grid was defined with the following parameters: grid box center at (6.375, 10.226, 9.454), number of points (70, 54, 52), and a grid spacing of 0.375 Å. This configuration encompassed the known peptide binding site, including residues R68, R69, P70, V71, A72, Q73, S74, G101, T102, Y120, V206, W207, G208, and Q209. The resulting docking poses were ranked by estimated Gibbs free energy and filtered to retain only molecules with molecular weights below 650 Da. For subsequent biophysical screening using SPR, 79 compounds with the lowest Gibbs free energy were selected.

## Accession Codes

The structure factors and atomic coordinates have been deposited in the Protein Data Bank under accession code 9RV5

## Author Contributions

C.L. prepared the figures and drafted the manuscript with contributions from F.F. and A.K. which was edited by K.S. and S.K.. C.L. developed and performed biophysical assays. F.L. and C.L. performed NMR measurements. C.L. developed and performed cellular assays with contributions from L.E.. A.K. and C.L. solved the crystal structure. J.D. performed *in silico* virtual screening. K.S., H.W., V.D., S.M.G., S.M., and S.K. supervised the research. This manuscript was edited and approved by all authors.

## Acknowledgments

The authors are grateful for support by the Structural Genomics Consortium (SGC), a registered charity (no. 1097737) that receives funds from Bayer AG, Boehringer Ingelheim, Bristol Myers Squibb, Genentech, Genome Canada through Ontario Genomics Institute [OGI-196], EU/EFPIA/OICR/McGill/KTH/Diamond Innovative Medicines Initiative 2 Joint Undertaking [EUbOPEN grant 875510], Janssen, Merck KGaA (aka EMD in Canada and U.S.), Pfizer, and Takeda. AK and SK would like to acknowledge funding from the German Cancer Consortium (DKTK) at the German Cancer Research Center (DKFZ). The Center for Biomolecular Magnetic Resonance (BMRZ) is supported by the state of Hesse. We thank the beamline scientist at Diamond Light Source for their support.

---

**Figure S1.**
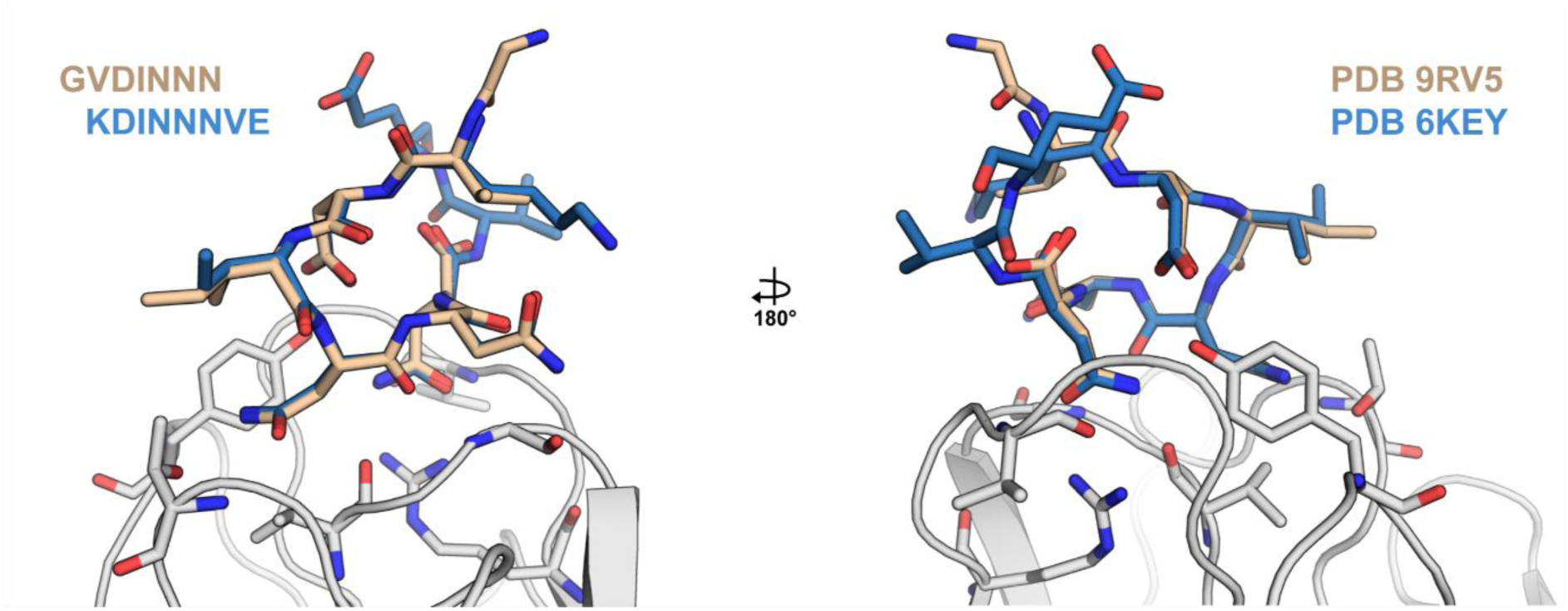
Superimposition of crystallized linear TAT peptide sequence GVDINNN in complex with hSPSB2^SPRY^ (PDB ID: 9RV5) and KDINNNVE (PDB ID: 6KEY)

**Figure S2.**
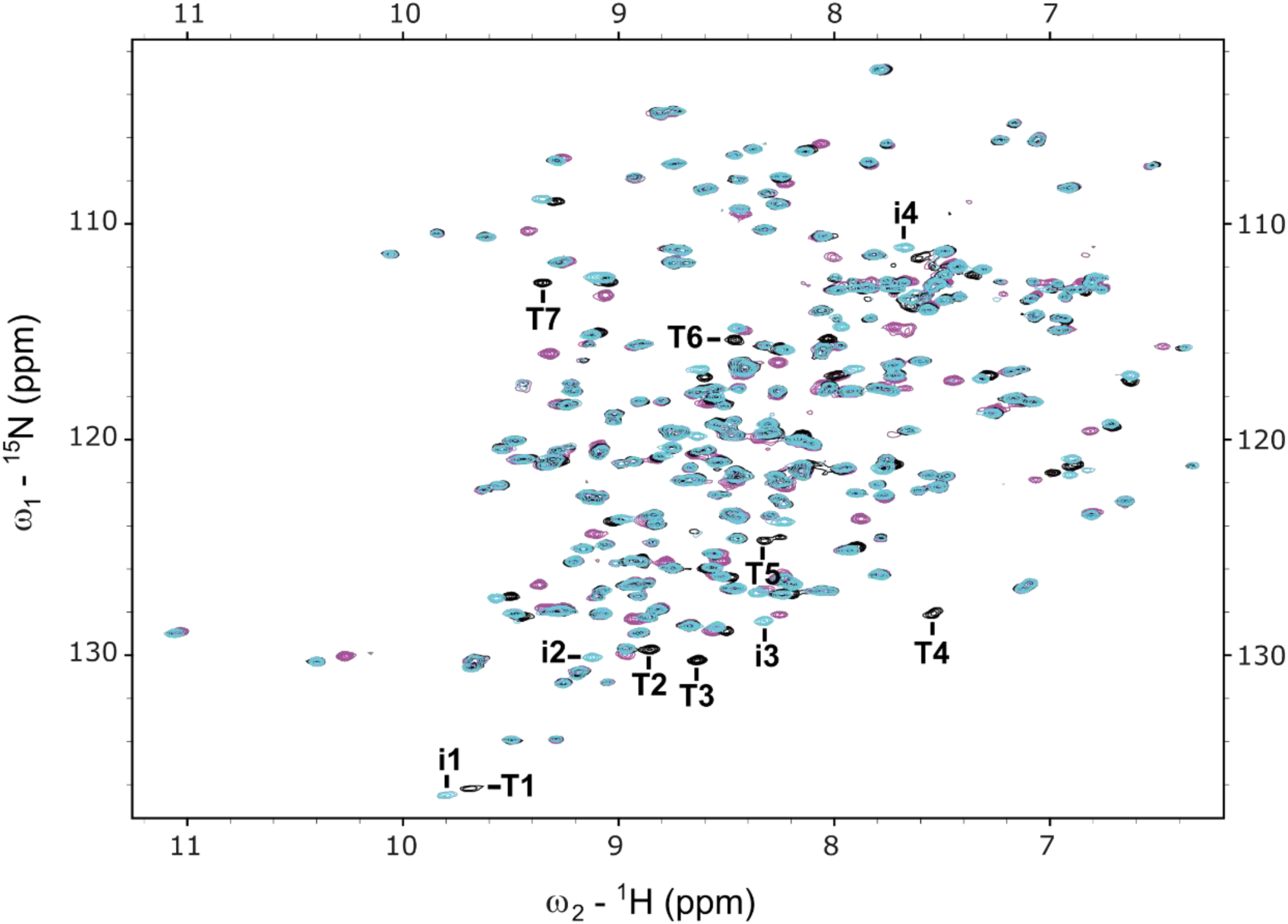
^1^H-^15^N BEST-TROSY spectra of ^15^N labelled hSPSB2^SPRY^ in the absence (magenta) and presence of linear TAT peptide (black) or iNOS peptide (cyan). Exemplary peaks that exhibit distinct chemical shift perturbations compared to the apo form and do not overlap are labelled as **i#** for the iNOS peptide and **T#** for the linear TAT peptide.

**Figure S3.**
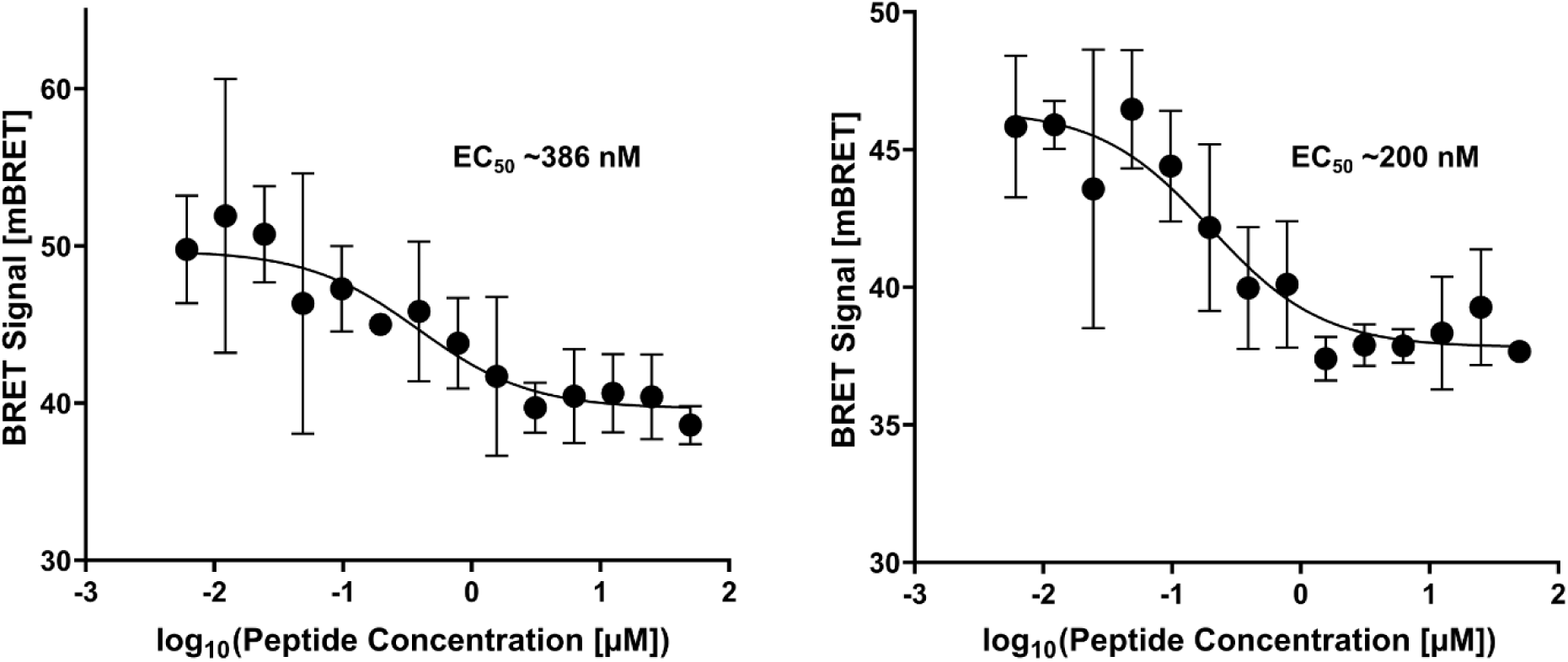
Displacement NanoBRET data of 30 nM cyclic TAT tracer (left) and 20 nM cyclic TAT tracer (right) with cyclic TAT peptide in intact HCT116 cells and respective EC50 values. Measured values are depicted as mean ± SD (n = 3).

**Supplement Table 1.** Overview of SPR single cycle kinetics results. KD, kon and koff values are depicted as mean ± SD (n = 3).

| Peptide | SPR $K_D$ [nM] | SPR $k_{on}$ [$\text{M}^{-1}\text{s}^{-1}$] | SPR $k_{off}$ [$\text{s}^{-1}$] |
| --- | --- | --- | --- |
| Linear RGD peptide | $162.0 \pm 4.0$ | $(2.0 \pm 0.1) \cdot 10^5$ | $(3.2 \pm 0.1) \cdot 10^{-2}$ |
| iNOS peptide | $0.8 \pm 0.1$ | $(2.9 \pm 0.1) \cdot 10^6$ | $(2.4 \pm 0.1) \cdot 10^{-3}$ |
| Linear TAT peptide | - | - | - |
| Cyclic TAT peptide | - | - | - |
| TAT motif | - | - | - |

**Supplement Table 2.**
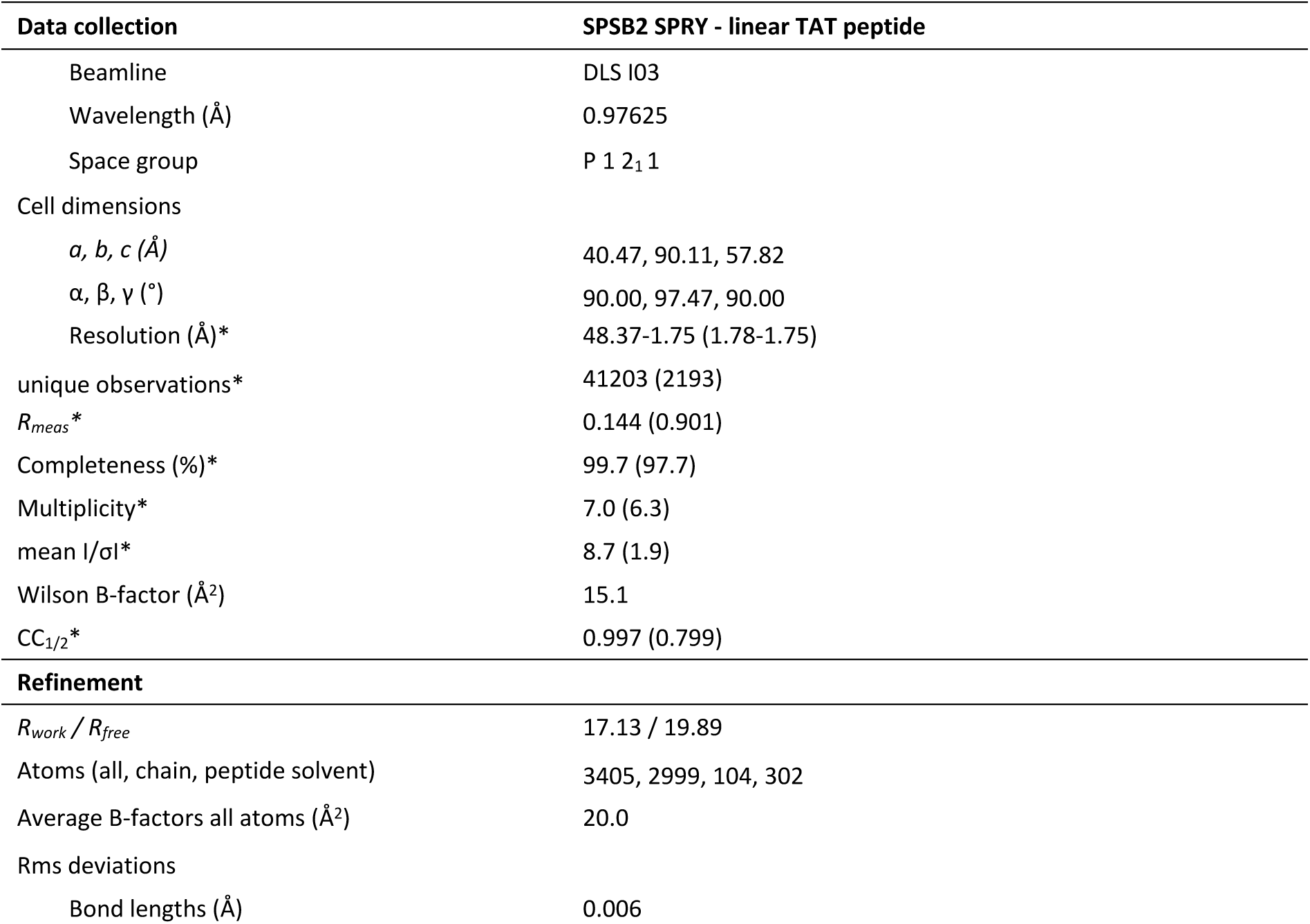
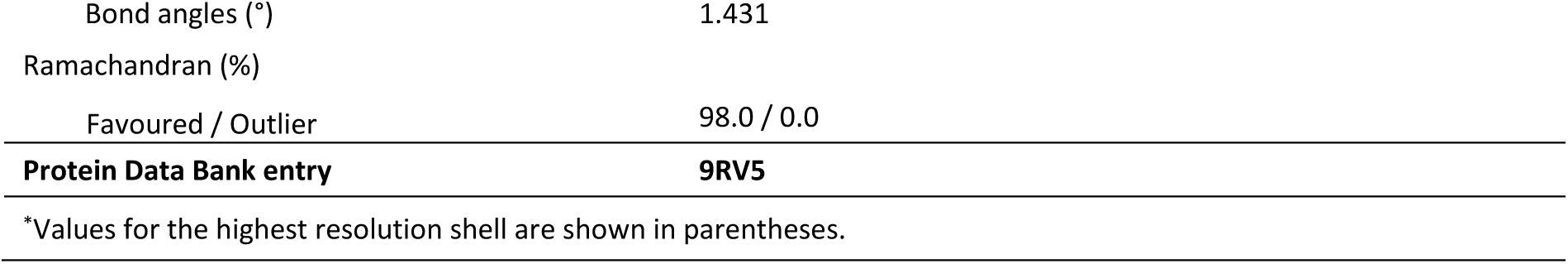
Data collection and refinement table.

## Notes

The authors declare no competing interest.

