## Supplemental Information for "Overcoming Ligand Discovery Challenges: Developing Peptide-Based Tracers for SPSB2"

**Supplement Table 1.** Overview of SPR single cycle kinetics results. K_D_, k_on_ and k_off_ values are depicted as mean ± SD (n = 3).

| **Peptide** | **SPR K_D_ [nM]** | **SPR k_on_ [M^-1^s^-1^]** | **SPR k_off_ [s^-1^]** |
| --- | --- | --- | --- |
| Linear RGD peptide | 162.0 ± 4.0 | (2.0 ± 0.1)*10^5^ | (3.2 ± 0.1)*10^-2^ |
| iNOS peptide | 0.8 ± 0.1 | (2.9 ± 0.1)*10^6^ | (2.4 ± 0.1)*10^-3^ |
| Linear TAT peptide | - | - | - |
| Cyclic TAT peptide | - | - | - |
| TAT motif | - | - | - |

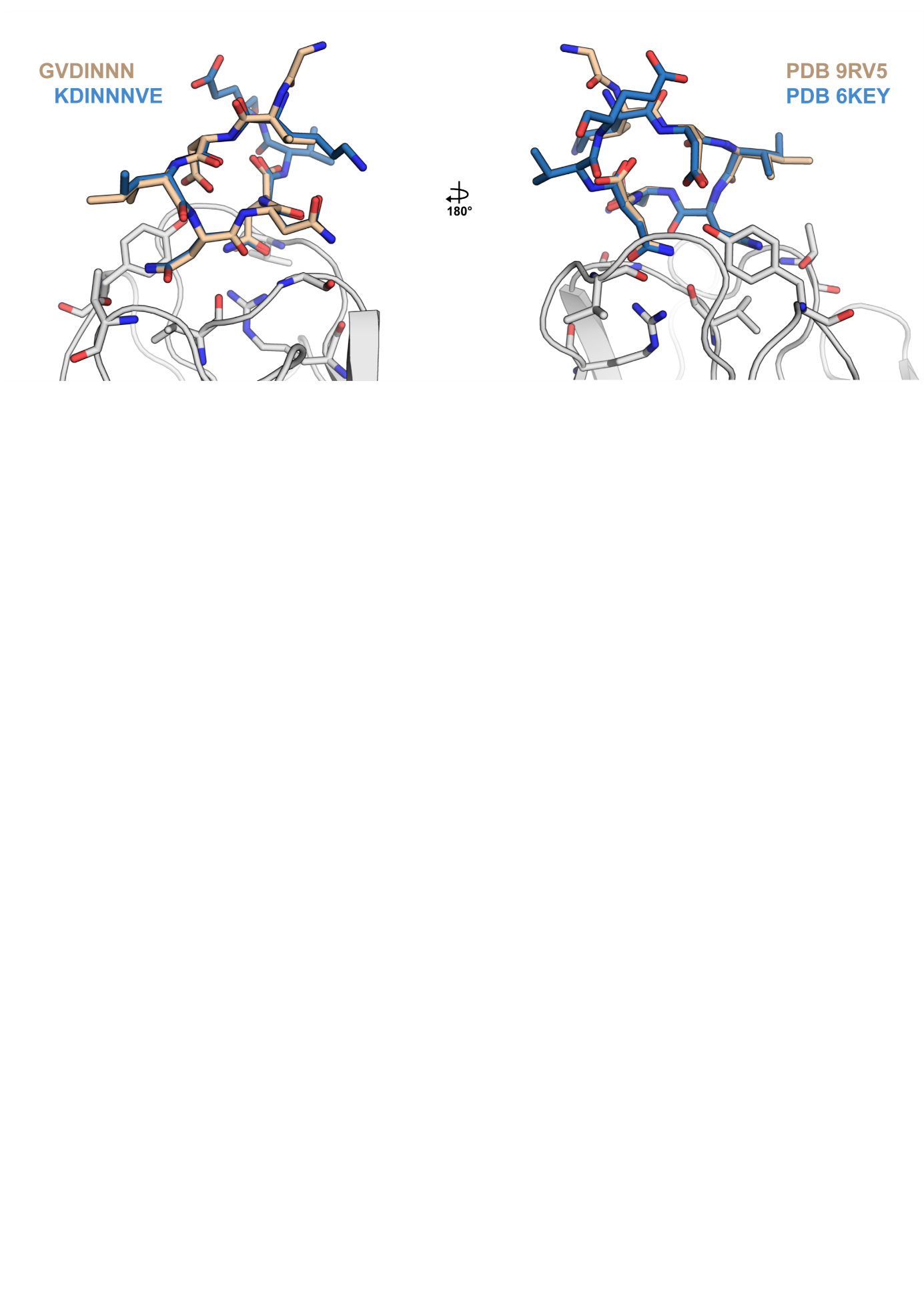

Figure S1 Superimposition of crystallized linear TAT peptide sequence GVDINNN in complex with hSPSB2^SPRY^ (PDB ID: 9RV5) and KDINNNVE (PDB ID: 6KEY)

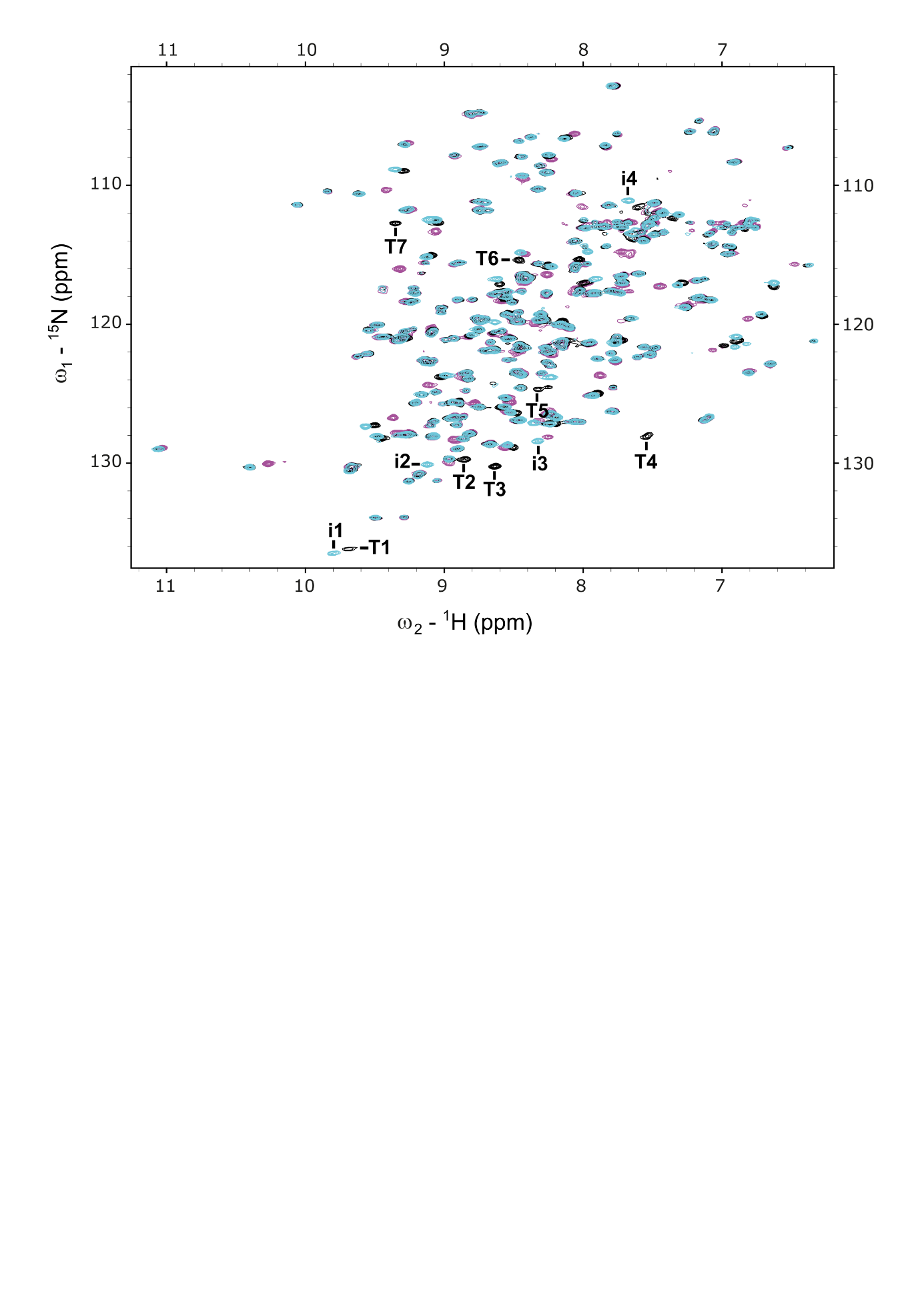

**Figure S2.** ^1^H-^15^N BEST-TROSY spectra of ^15^N labelled hSPSB2^SPRY^ in the absence (magenta) and presence of linear TAT peptide (black) or iNOS peptide (cyan). Exemplary peaks that exhibit distinct chemical shift perturbations compared to the apo form and do not overlap are labelled as **i#** for the iNOS peptide and **T#** for the linear TAT peptide.

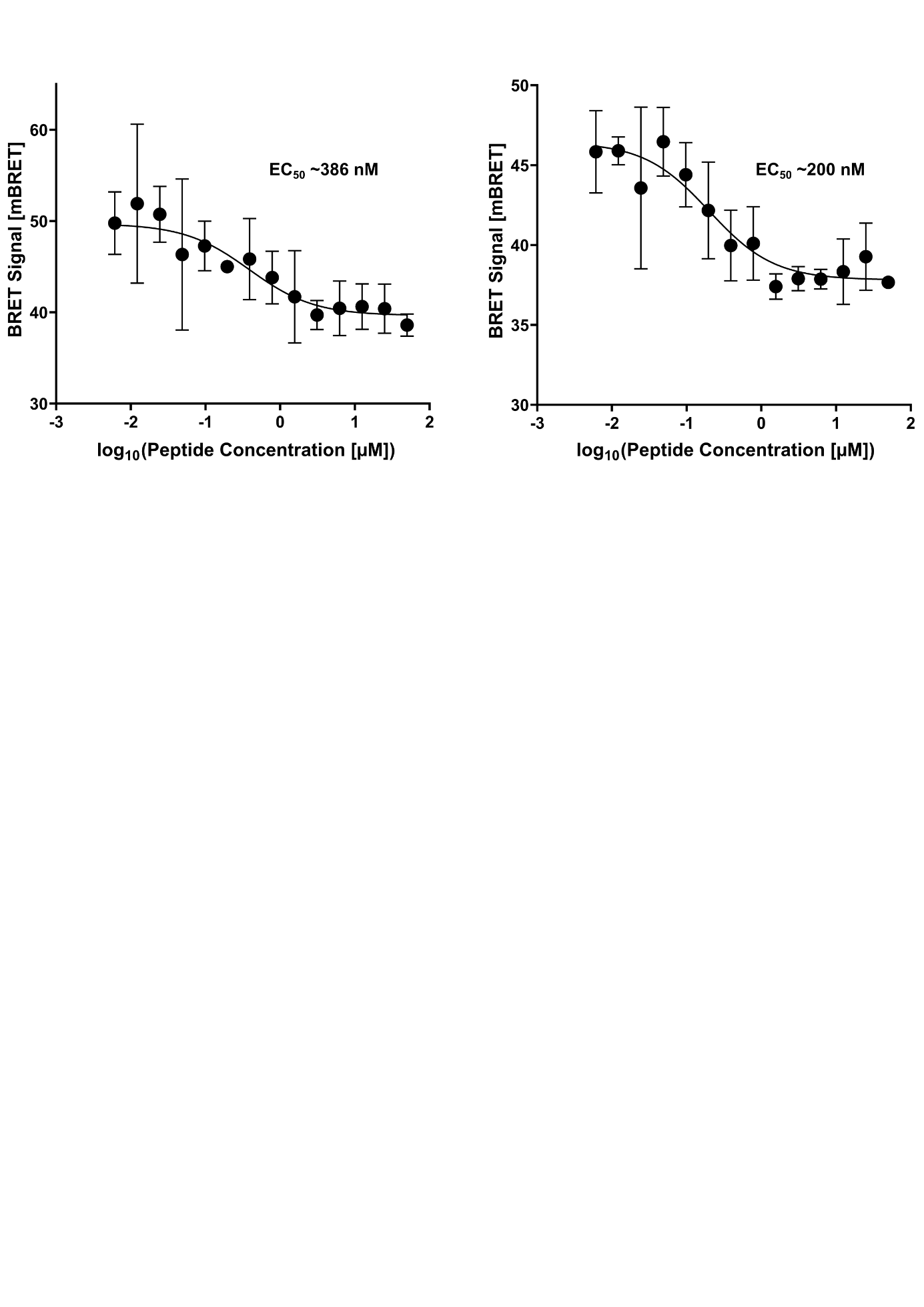

**Figure S3.** Displacement NanoBRET data of 30 nM cyclic TAT tracer (left) and 20 nM cyclic TAT tracer (right) with cyclic TAT peptide in intact HCT116 cells with respective EC_50_ values. Measured values are depicted as mean ± SD (n = 3).

**Supplement Table 2.** Data collection and refinement table

| **Data collection** | **SPSB2 SPRY - linear TAT peptide** |
| --- | --- |
| Beamline | DLS I03 |
| Wavelength (Å) | 0.97625 |
| Space group | P 1 2_1_ 1 |
| Cell dimensions |  |
| *a, b, c (Å)* | 40.47, 90.11, 57.82 |
| α, β, γ (°) | 90.00, 97.47, 90.00 |
| Resolution (Å)* | 48.37-1.75 (1.78-1.75) |
| unique observations* | 41203 (2193) |
| *R_meas_** | 0.144 (0.901) |
| Completeness (%)* | 99.7 (97.7) |
| Multiplicity* | 7.0 (6.3) |
| mean I/σI* | 8.7 (1.9) |
| Wilson B-factor (Å^2^) | 15.1 |
| CC_1/2_* | 0.997 (0.799) |
| **Refinement** |  |
| *R_work_ / R_free_* | 17.13 / 19.89 |
| Atoms (all, chain, peptide solvent) | 3405, 2999, 104, 302 |
| Average B‐factors all atoms (Å^2^) | 20.0 |
| Rms deviations |  |
| Bond lengths (Å) | 0.006 |
| Bond angles (°) | 1.431 |
| Ramachandran (%) |  |
| Favoured / Outlier | 98.0 / 0.0 |
| **Protein Data Bank entry** | **9RV5** |
| ^*^Values for the highest resolution shell are shown in parentheses. | |
